# Live *Malassezia* strains isolated from the mucosa of patients with ulcerative colitis

**DOI:** 10.1101/2023.11.28.569113

**Authors:** Yong-Joon Cho, Juan Yang, Seung Yong Shin, Hyo Keun Kim, Piyapat Rintarhat, Minji Park, Muhyeon Sung, Kate Lagree, David M. Underhill, Dong-Woo Lee, Chang Hwan Choi, Chul-Su Yang, Won Hee Jung

## Abstract

The human gut is inhabited by a complex ecosystem of diverse microorganisms. While most studies on the gut microbiota have focused on bacteria, accumulating evidence has underscored the role of the mycobiota in inflammatory bowel disease (IBD). This study is the first to isolate and characterize live *Malassezia globosa* strains from the intestinal mucosa of patients with ulcerative colitis. *Malassezia* species primarily inhabit the human skin. We therefore compared the *M. globosa* gut isolates with the *M. globosa* skin isolates and noted a striking disparity between them. The gut isolates led to a greater exacerbation of colitis in mice. Transcriptome analysis revealed that the gut isolates were more sensitive to normoxia than the skin isolates, suggesting adaptation to hypoxia prevalent in the intestinal environment. These findings provide novel insights into the potential impact of *M. globosa* on the pathogenesis of IBD and the influence of niche-specific adaptations on its virulence.

## Introduction

Inflammatory bowel disease (IBD) is a chronic and relapsing–remitting disease that poses a considerable healthcare burden globally. The prevalence of IBD has significantly increased not only in Europe and North America but also in East Asian countries, including Korea. Numerous studies have demonstrated that IBD is caused by various factors, including the genetic susceptibility of the host, dysbiosis of the microbiome, disruption of intestinal barrier functions, and inappropriate immune activation, resulting in the disruption of intestinal homeostasis ^1,2^. Ulcerative colitis (UC) and Crohn’s disease (CD) are the two types of IBD. UC occurs as a continuous superficial inflammation mostly in the mucosa and submucosa of the colon, while CD presents as scattered lesions throughout the gastrointestinal tract ^3^. In general, Western countries, such as those in North America, Western Europe, and Oceania, have a higher prevalence of CD than UC ^4,5^. Although the total number of IBD cases has increased, the number of UC cases and their annual increase are significantly higher (approximately more than twofold) in Korea ^6^. The pathogenesis of UC and CD are still not fully understood; however, similar factors influence both types of IBD ^7^.

The microbiome is a crucial factor in the intestinal environment. In general, the intestinal microbiome in the mammalian host is well controlled by multiple mechanisms, such as the production of antimicrobial peptides and secretion of immunoglobulin A (IgA) and IgG, which can prevent epithelial invasion by microbes. Moreover, mucin, which is a dense hydrogel layer, covers the surface of intestinal epithelial cells and is produced by goblet cells within the epithelium ^8^. However, numerous studies have reported a shift in the intestinal microbiome in patients with UC and CD ^9,10^. Although significant individual variation has been noted, numerous studies have reported common features of alteration of the intestinal microbiome, including reduced bacterial diversity, in patients with IBD. In particular, decreased abundance of Firmicutes phylum and the association of proinflammatory bacteria, such as *Escherichia coli*, *Ruminococcus gnavus*, and *Fusobacterium* species, with IBD have been reported ^11^.

In addition to the bacterial community, the fungal microbiome (known as the mycobiome) likely plays a crucial role in the gut. Notably, although metagenomic data suggest that fungal DNA accounts for less than 0.1% of the intestinal microbiome, the fungal community must be considered because of the size of fungal cells, which is usually more than 100-fold larger than that of bacterial cells, when studying the intestinal microbiome ^12^. Accumulating evidence indicates that the mycobiome contributes to the development of IBD, and several fungi commonly observed in multiple studies are associated with individual susceptibility to IBD ^12^. A pioneering study revealed that the increased abundance of fungal populations, particularly *Candida* species, plays an essential role in antifungal innate immunity in a dextran sodium sulfate (DSS)-induced colitis mouse model ^13^. *Candida* species are most frequently found in the human intestinal mycobiota and have long been considered to be commensal fungi in the human gut. Although the role of *Candida* species in IBD remains controversial, a clinical study revealed that the abundance of *Candida* species in the intestinal mycobiota of patients with UC is positively correlated with favorable clinical outcomes of fecal transplantation ^14^. Moreover, numerous studies have demonstrated the augmentation of colitis by oral gavage of *Candida* species in a DSS-induced colitis mouse model ^12^.

*Malassezia* is considered the predominant fungus in the human skin. It is a pathobiont involved in various skin diseases, including psoriasis, seborrheic dermatitis, and atopic dermatitis ^15^. Studies have indicated a possible role of *Malassezia* species in chronic diseases occurring at other body sites, such as pancreatic cancer and Alzheimer’s disease ^16–18^. Recent mycobiome analyses have frequently observed *Malassezia* DNA sequence reads in the human gut indicating that the fungus is one of the core taxa in the human intestinal mycobiota ^19–21^. A positive correlation between *Malassezia* and *Bacteroides*, which is a predominant bacterial genus in the human gut, has been reported ^22^. Furthermore, studies utilizing skin swabs and fecal samples from infants suggested that *Malassezia* is transmitted from the mother and colonized in the gut of infants, although the abundance of the fungus gradually decreases upon aging ^23,24^. These findings suggest that *Malassezia* resides in both skin and gut mucosa and that the fungus has evolutionally adapted to a distinct niche-specific environment.

To date, 18 different species have been identified within the *Malassezia* genus ^15^. Among these, a recent study by Limon et al. using mucosa-washing samples revealed that *M. restricta* and *M. globosa* were strongly associated with CD ^25^. In that study, a higher abundance of *Malassezia* was noted in the intestinal mucosal surface of patients with CD than in that of healthy individuals. Moreover, in the same study, patients with CD with the IBD risk allele caspase recruitment domain-containing protein (CARD)^S12N^ variant exhibited a tight association with the presence of *Malassezia* in their intestinal mucosa, further confirming the positive correlation between the fungus and the disease ^25^. CARD9 plays a crucial role in a central signaling pathway in innate immune responses and is activated by pattern recognition receptors, including C-type lectin receptors, such as Dectin-1, Dectin-2, and Mincle, which are key receptors for detecting commensal and pathogenic fungi. Moreover, CARD9^S12N^ is a crucial single-nucleotide polymorphism (SNP) involved in IBD ^26^. *Malassezia* also exacerbated colitis in a DSS-induced colitis mouse model, which required functional CARD9. Furthermore, Dectin-2, but not Dectin-1 and Mincle, played a crucial role in responding to *Malassezia* in the mouse model ^25^. Thus, the findings of Limon et al. provide insights into the possible contribution of *Malassezia* to IBD. However, in their study, Limon et al. used *M. restricta* MYA-4611, synonymous with the *M. restricta* type strain CBS 7877, which may not properly represent the fungus within the gut environment because it was originally isolated from the human skin and not from the human gut ^27^. In general, niche-specific metabolic adaptation, which is normally linked to genomic variations and niche-dependent gene expression, is a well-established phenomenon, particularly in bacteria, based on phenotypic and multiomics analysis ^28–30^. Fungi also reside in various environments, including different niches within the human host, and multiple selective pressures, such as oxygen depletion, nutrient limitation, and pH fluctuation, lead to niche-specific adaptation ^31,32^. To date, no comprehensive study has assessed phenotypic and genetic diversities in the *Malassezia* population in different host niches within the human body to demonstrate niche-specific adaptation of the fungus at the strain level.

In this study, we hypothesized that *Malassezia* adapts to different host niches, such as the skin and gut, and therefore possesses different phenotypic characteristics, including virulence, at the strain level. To confirm our hypothesis, we isolated a *Malassezia* strain directly from the human intestinal mucosal surface of patients with UC and assessed its genome and virulence in comparison with those of the same fungal species isolated from the human skin. We successfully isolated live *M. globosa* strains from the intestinal mucosa of patients with UC. We compared the genomes, transcriptomes, and virulence of the *M. globosa* gut isolates with those of the *M. globosa* skin isolates to identify differences in genotypes and phenotypes caused by potential niche-specific adaptations of the fungus.

## Results

### Mycobiota analysis identified that *Malassezia* resides in the mucosal layer of the human intestine in patients with UC

A study demonstrated that microbiomes in murine and human fecal samples only partially represent gut mucosal microbiomes ^33^. Moreover, previous mycobiome studies suggested that numerous fungi detected in fecal samples are temporal and originate from the environment, making it difficult to differentiate the true fungal colonizers in the gut ^34,35^. Alternatively, analysis of mucosal layer-associated fungal communities provided more reliable data and revealed a strong correlation between the gut mycobiota and IBD, such as CD ^25^. Therefore, in this study, we first compared the mycobiota associated with the intestinal mucosal surface of patients with UC and healthy individuals by collecting water lavage samples during colonoscopy. Furthermore, to determine whether inflammation influences the composition of the mycobiota associated with the intestinal mucosal layer, we collected water lavage samples from the intestinal mucosal surfaces with and without inflammation from the same patient with UC. In addition, as mentioned earlier, we incubated a portion of each water lavage sample on a *Malassezia*-specific medium to isolate live *Malassezia* cells from the human gut mucosa.

In total, 56 intestinal water lavage samples were obtained from 29 patients during colonoscopy. Of these, 54 were paired (one from the intestinal mucosal surface with inflammation and the other without inflammation from the same patient). Two samples were obtained only from the intestinal mucosal surface without inflammation from patients with UC. Moreover, 11 water lavage samples were obtained from 11 healthy individuals and included in this study. Detailed information on the samples is provided in Table 1 and Supplementary Table S1. Once the water lavage samples (approximately 50 mL) were obtained, half of the samples were immediately inoculated on Leeming and Notman agar (LNA) medium within three hours without freezing to minimize the loss of live fungal cells and incubated in the presence of 2% oxygen to mimic the intestinal environmental condition. The remaining samples were subjected to DNA extraction for amplicon sequencing. In total, 6,913,621 reads were obtained from 67 samples. After quality filtering and chimera removal, 6,621,891 reads were mapped to 1,074 amplicon sequence variants. *Nakaseomyces*, *Candida*, *Saccharomyces*, and *Aspergillus* were the top four abundant genera throughout the samples (Fig. 1A). The fifth most abundant fungal genus in all samples was *Malassezia*.

**Fig. 1.**
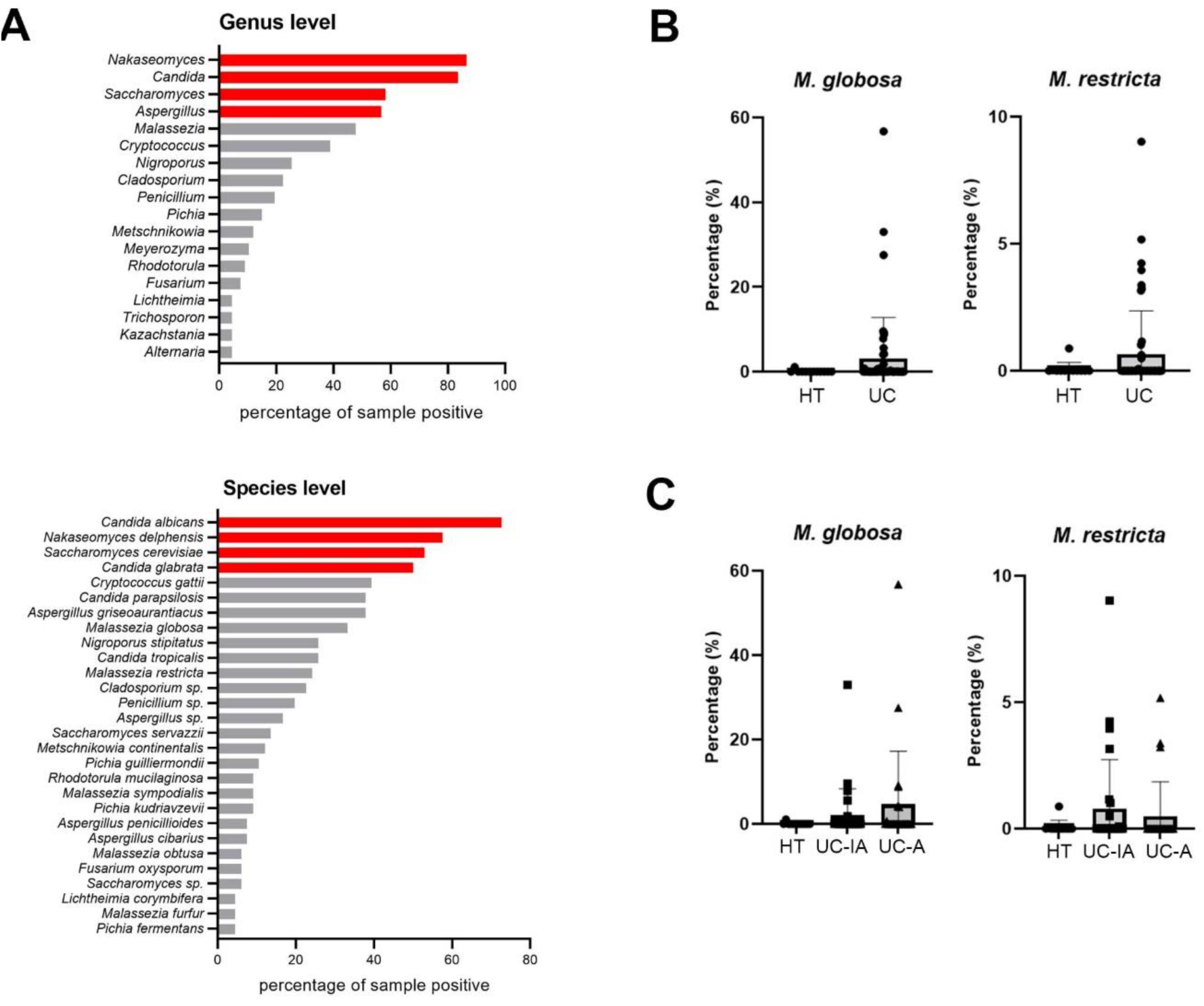
Results of amplicon sequencing analysis. **A.** Analysis of fungal genera and species most frequently present in all 67 samples analyzed in this study. Fungal genera and species depicted in red bars are present in more than 50% of each sample. **B.** The rank test revealed that more *M. globosa* and *M. restricta* reads were present in UC samples than in healthy control samples (HT); however, the significance of the difference was low. **C.** The rank test revealed no significant difference in the abundance of *M. globosa* and *M. restricta* between the gut mucosal surfaces with (UC-T) and without (UC-F) inflammation in the same patient with UC.

**Table 1.** Demographic and clinical characteristics of the participants.

| Characteristics | Patients with UC<sup>a)</sup> | Healthy controls |
| --- | --- | --- |
| Number | 29 | 11 |
| Age (years) | 48.5 ± 17.3 | 61.3 ± 9.9 |
| Male sex, No. (%) | 17 (58.6) | 4 (36.4) |
| Disease extent (maximum), No. (%) |  |  |
| Proctitis | 2 (6.9) |  |
| Left-sided colitis | 8 (27.6) |  |
| Extensive colitis | 19 (65.5) |  |
| Endoscopic subscore, No. (%) |  |  |
| 0 | 3 (10.4) |  |
| 1 | 7 (24.1) |  |
| 2 | 16 (55.2) |  |
| 3 | 3 (10.3) |  |
| Sampling site, No (%) |  |  |
| With inflammation |  |  |
| Rectum | 4 (13.8) |  |
| Sigmoid colon | 15 (51.7) |  |
| Descending colon | 5 (17.2) |  |
| Transverse colon | 0 (0.0) |  |
| Ascending colon | 2 (7.0) |  |
| None | 3 (10.3) |  |
| Without inflammation |  |  |
| Rectum | 4 (13.8) | 0 (0.0) |
| Sigmoid colon | 5 (17.2) | 4 (36.3) |
| Descending colon | 10 (34.5) | 5 (45.5) |
| Transverse colon | 8 (27.6) | 0 (0.0) |
| Ascending colon | 2 (6.9) | 2 (18.2) |
| Concomitant medication (Overlapped), No. (%) |  |  |
| Immunomodulators | 21 (72.4) |  |
| Prior use of biological agents | 16 (55.2) |  |
| One medication | 10 (62.5) |  |
| Two medications and above | 6 (37.5) |  |
| Prior use of JAK inhibitors<sup>b)</sup> | 2 (6.9) |  |

<sup>a)</sup>UC, Ulcerative colitis; <sup>b)</sup>JAK inhibitors, Janus kinase inhibitors

We performed α- and β-diversity analyses using the samples. Unlike our expectation, no significant differences were noted between patients with UC and healthy individuals or between the sites with inflammation and those without inflammation at the intestinal mucosal surface of patients with UC (Supplementary Fig. S1). A rank test was performed to determine whether the abundance of *Malassezia* is associated with UC. The abundance of *M. globosa* and *M. restricta* tended to be higher in patients with UC than in healthy individuals; however, the statistical significance of the difference was low (Fig. 1B). Moreover, no significant difference was noted in the abundance of both *M. globosa* and *M. restricta* between the intestinal mucosal surfaces with and without inflammation in each patient with UC, indicating a lack of correlation between the presence of inflammation and the abundance of *Malassezia* (Fig. 1C). Overall, although our data suggested no significant difference in the diversity of the mycobiota between patients with UC and healthy individuals and between mucosal samples with and without inflammation from the same patient, patients with UC tended to have a higher abundance of *M. globosa* and/or *M. restricta* in their gut mucosal surface than healthy individuals.

### Isolation of *M. globosa* from the intestinal mucosa and comparison with *M. globosa* skin isolates

In addition to performing mycobiota analysis by amplicon sequencing, water lavage samples were used to isolate live *Malassezia* strains from the intestinal mucosa. Live fungal strains belonging to 28 and seven different species were isolated from patients with UC and healthy individuals, respectively (Supplementary Table S2). Among these, two *M. globosa* strains were identified: one was isolated from the sigmoid colon without inflammation from patient no. 21 with UC and the other was isolated from the same site but with inflammation from patient no. 50 with UC (Table S1). These two *M. globosa* strains were deposited at the Korean Collection for Type Cultures (KCTC; https://kctc.kribb.re.kr/en) and designated KCTC 37188 and KCTC 37189, respectively. Notably, both patients with UC displayed *M. globosa* reads in the amplicon sequencing data for mycobiota analysis.

To determine whether *M. globosa* strains isolated from two different host niches, namely the gut mucosal surface and the skin, possess distinct characteristics, the genomes of the gut isolates *M. globosa* KCTC 37188 and KCTC 37189 and the skin isolates *M. globosa* KCTC 27541 and KCTC 27776, which were obtained from the forehead of patients with seborrheic dermatitis in our previous study ^36^, were sequenced using Illumina sequencing technology. Average nucleotide identity (ANI) was calculated based on the genome sequences of each strain and *M. globosa* CBS 7966, a type strain of the species that was isolated from the skin of a patient with pityriasis versicolor in England (https://wi.knaw.nl) ^37^. Overall, the ANI values between the gut and skin isolates were relatively high, indicating that the genome sequences of each strain were highly similar and exhibited no clear difference (Fig. 2A). Moreover, relatively low ANI values were observed between the *M. globosa* CBS 7966 type strain and the gut and skin isolates, possibly because of geological and ethnic differences in the isolation sites of the strains. A high similarity between the genomes of the gut and skin isolates was also confirmed by SNP and indel analyses on comparing the genomes with the reference genome of the type strain (Fig. 2B). Furthermore, homologous clustering revealed that the gut and skin isolates and the type strain shared 3,820 common homologs (Fig. 2C). Few unique genes were identified in each strain: 30, 15, 20, 14, and 26 in the genomes of strains KCTC 37188, KCTC 37189 (gut isolates), KCTC 27541, KCTC 27777 (skin isolates), and CBS 7966, respectively. However, most unique genes in each strain were hypothetical and were predicted to produce a protein with an unknown function.

**Fig. 2.**
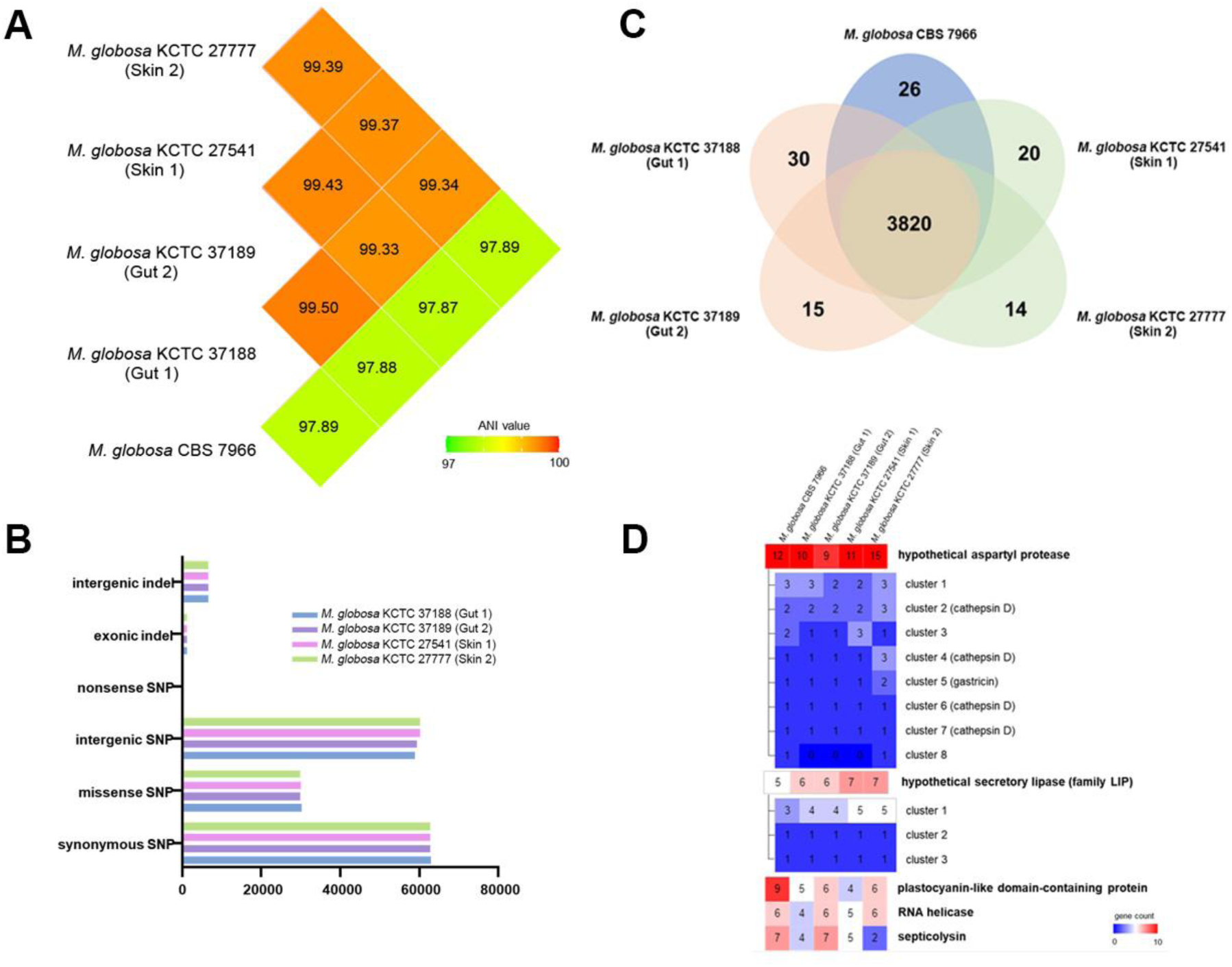
Genome comparison of *M. globosa* strains isolated from the gut and skin. **A.** Comparison of ANI values between *M. globosa* strains. **B.** A bar graph representing the results of SNP and indel analyses. **C.** A phi chart showing the results of homologous clustering analysis, indicating the number of shared and unique genes in each strain. **D.** A list of cluster groups, where the number of genes belonging to each orthologous cluster group differs among strains. The top five groups with the highest average number of genes were selected.

Previous studies on genomes of multiple *Malassezia* species have suggested that *Malassezia* contains many genes that have high copy numbers. Most of these genes were predicted to contribute to the virulence of *Malassezia* and may be crucial for the survival of the fungus in the host environment. Examples include genes encoding lipases and proteases ^38^. In this study, we also analyzed genes with multiple copy numbers in the genomes of the gut and skin isolates and the type strain to identify the enrichment of any gene family in a specific isolate. The copy numbers of genes encoding protease and lipase were highly enriched across the strains, and no particular feature was observed (Fig. 2D). Moreover, we paid particular attention to the copy number of the gene encoding the homolog of septicolysin, which may be horizontally transferred from a bacterium ^39^, because of its putative endotoxin activity. The copy number of the septicolysin homolog gene varied across the strains; it was dependent on individual strains and not on the isolation site of each strain. Overall, no specific genomic characteristics were noted in the *M. globosa* gut isolates, especially in terms of nucleotide sequences and gene content levels, compared with the skin isolates and the type strain in this study.

### *M. globosa* isolated from the intestinal mucosal surface is more sensitive to oxygen than that isolated from the skin

In addition to the genome contents, we compared the transcriptome profiles between the gut and skin isolates. We used different oxygen concentrations because oxygen plays a crucial role in the intestinal environment ^40,41^ and hypothesized that the transcriptome profiles of the gut and skin isolates respond distinctly to different concentrations of oxygen. To compare the transcriptomes, we grew *M. globosa* gut and skin isolates in the presence of 2% and 20% oxygen, which likely represent the oxygen concentrations at the gut mucosal surface and skin surface, respectively, and analyzed and compared the transcriptome profile of each strain using RNA sequencing (RNA-Seq). Two independent gut isolates, KCTC 37188 and KCTC 37189, showed a relatively higher number of differentially expressed genes (83 and 108, respectively) than the skin isolates KCTC 27541 and KCTC 27777 (55 and 44, respectively) in response to increased oxygen concentrations (Fig. 3A). We next conducted an enrichment analysis of KEGG pathways and GO terms of differentially regulated genes in each strain under different oxygen concentrations. In the gut isolates grown in the presence of 20% oxygen, highly enriched KEGG pathways and GO terms representing genes involved in peroxisome and ARF protein signal transduction were detected (Fig. 3B). As one of the major functions of peroxisomes is scavenging cells from oxidative stress, these results indicated the *M. globosa* gut isolates were under oxidative stress in the presence of 20% oxygen ^42,43^. Furthermore, the homologs in the ARF protein signal transduction are involved in the regulation of stress responses via unfolded protein responses in *S. cerevisiae*, supporting our finding that the *M. globosa* gut isolates were under stress in the presence of high concentrations of oxygen ^44^. In contrast, we did not note the enrichment of any specific gene family in the *M. globosa* skin isolates grown in the presence of 20% oxygen. When the *M. globosa* gut isolates were grown in the presence of 2% oxygen, genes involved in translation and ribosome functions were highly enriched, suggesting that the gut isolates were more actively growing at lower oxygen concentrations than the skin isolates. These results suggest that the *M. globosa* gut isolates are more sensitive to a higher oxygen concentration and exhibit better fitness at a lower oxygen concentration than the skin isolates.

**Fig. 3.**
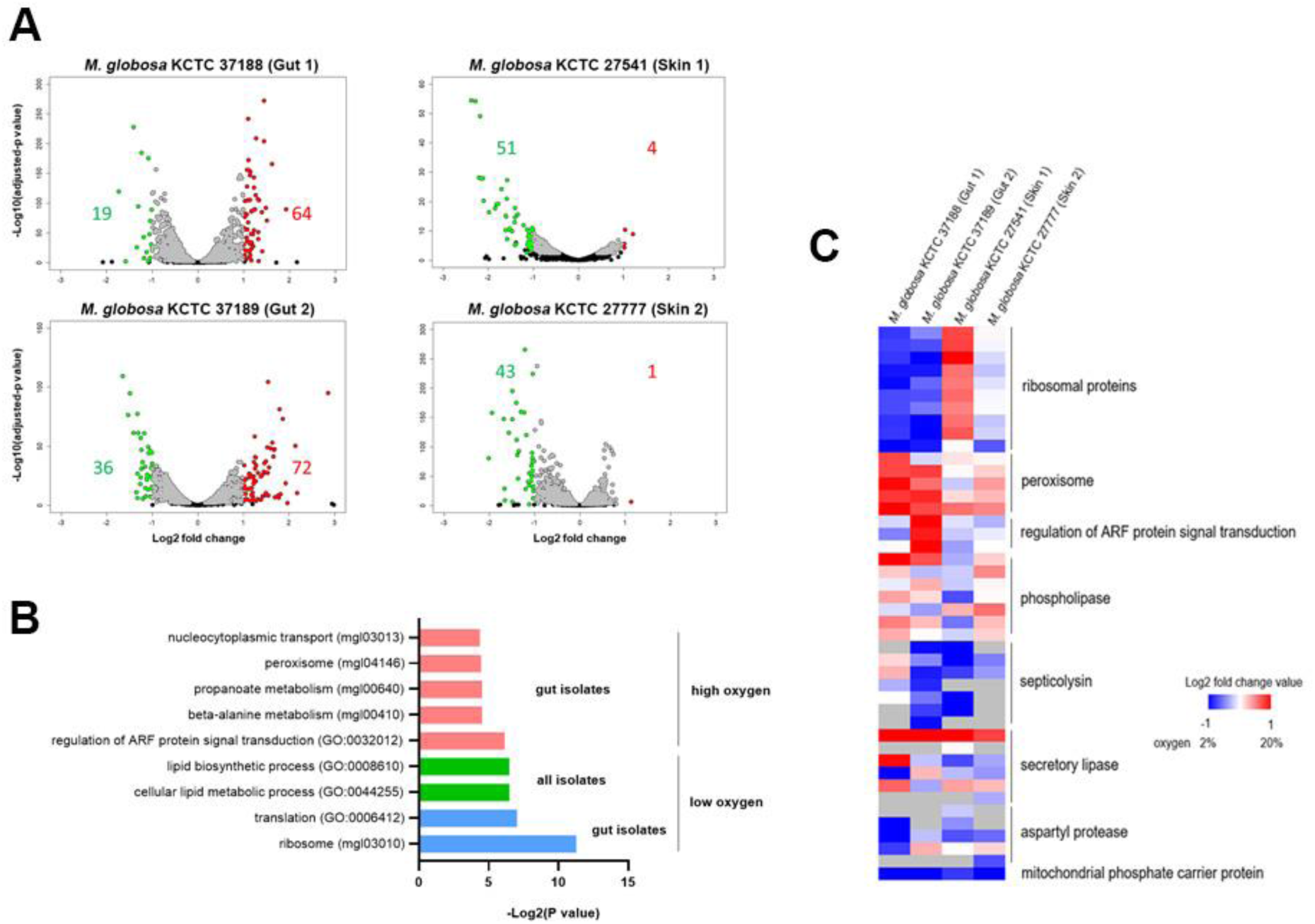
Comparison of transcriptomes of *M. globosa* strains isolated from the gut and skin. **A.** Strains were grown under low-oxygen (2%) or high-oxygen (20%) conditions, and transcriptome profiles were analyzed by RNA-Seq. Volcano plots show the differential gene expression in fungal cells grown under high-oxygen conditions in comparison with those grown under low-oxygen conditions. Red: upregulated genes under low-oxygen conditions. Green: the number of downregulated genes under low-oxygen conditions. **B**. Bar plots show the enrichment of GO terms and KEGG pathways in differentially expressed genes in each strain. Red: upregulated genes from gut-derived strains under high-oxygen conditions. Green: upregulated genes from both gut- and skin-derived strains under low-oxygen conditions. Blue: upregulated genes from gut-derived strains under low-oxygen conditions. **C**. A heatmap of selected gene clusters from transcriptome analysis.

Individual gene families that were specifically differentially regulated in the *M. globosa* gut isolates under the 20% oxygen condition compared with the 2% oxygen condition also included genes associated with ribosomal proteins and peroxisome pathways (Fig. 3C). In particular, the genes encoding ribosomal proteins were significantly downregulated in the gut isolates under the 20% oxygen condition, further confirming a decrease in protein synthesis and a resulting decrease in growth rate at a higher oxygen concentration. Moreover, the genes associated with the peroxisome pathway were highly upregulated in the gut isolates, also suggesting that the fungal cells are under oxidative stress and confirming the results of our KEGG and GO term enrichment analyses. Overall, transcriptome analysis suggested that the *M. globosa* gut isolates respond to oxygen concentrations differently from the skin isolates. Moreover, the results of transcriptome analysis suggested that the *M. globosa* gut isolates have evolved to adapt to the low-oxygen condition in the gut environment but are more sensitive to higher oxygen concentrations than the skin isolates.

### *M. globosa* isolated from the gut mucosa exacerbates DSS-induced colitis in mice

We next investigated whether the *M. globosa* gut isolates are associated with exacerbating colitis in the mouse model and compared their pathogenesis with those of the skin isolates. We treated the mice with DSS for 5 days to induce colitis. After DSS treatment, we inoculated the fungal cells of each strain by oral gavage every other day for a total of three times and monitored the disease progress (Fig. 4A). We also compared the virulence and contribution of the gut isolates to colitis with those of the skin isolates to determine whether strains from different host niches influence the disease.

**Fig. 4.**
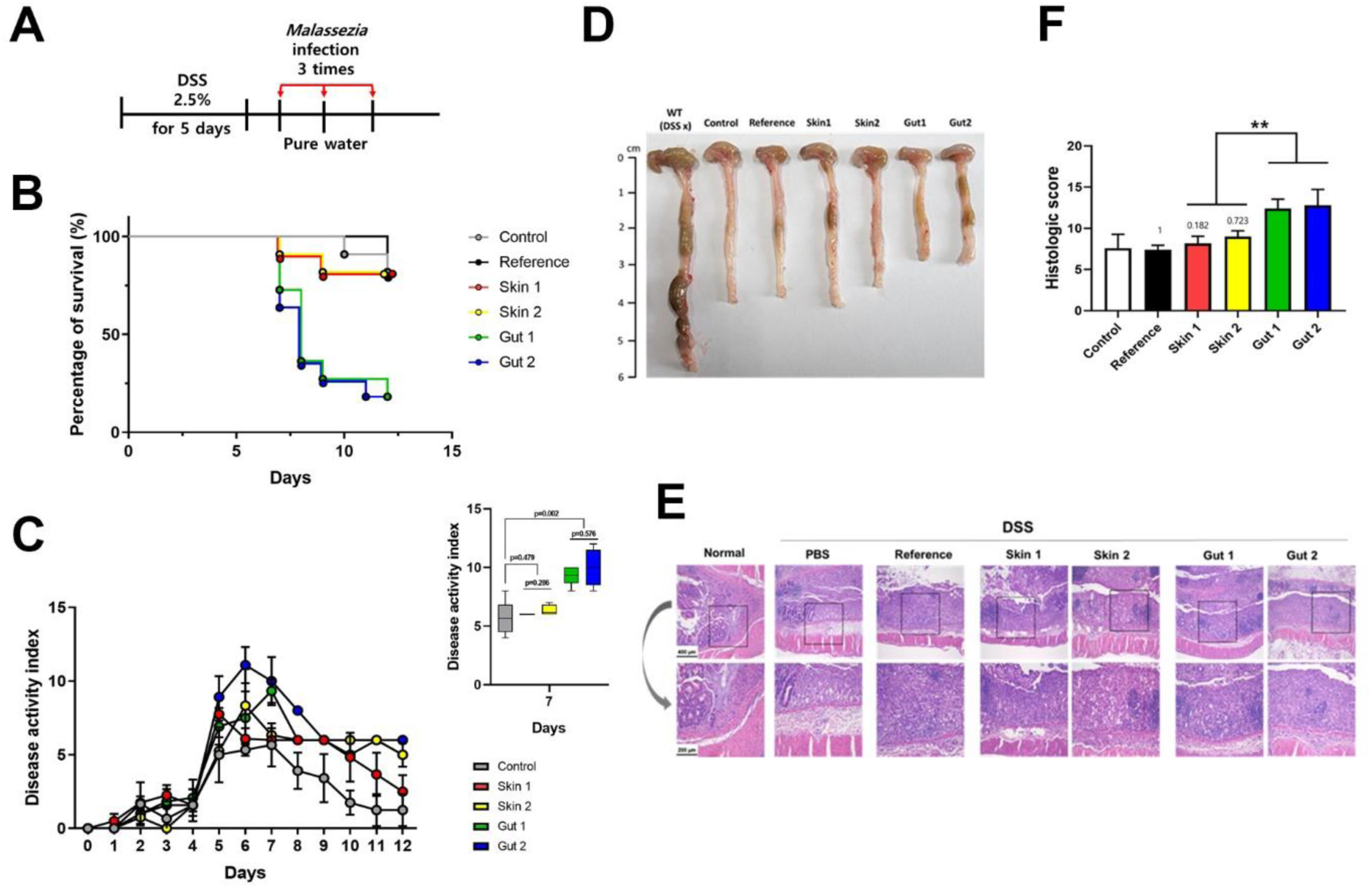
*M. globosa* isolated from the gut mucosa exacerbated DSS-induced colitis in mice. **A**. Colitis was induced by DSS treatment for 5 days. *M. globosa* strains were orally gavaged a total of three times every other day, and the survival of mice and disease progression were monitored. **B.** The survival of mice gavaged with each *M. globosa* strain was monitored. For each strain, 11 mice were infected. However, the reference strain *S. cerevisiae* ATCC 204508 was used to infect five mice. **C.** DAI scores of mice gavaged with the indicated fungal strains. A small box plot presents significantly different DAI scores on day seven. **D**. Representative colons from mice with DSS-induced colitis gavaged with the indicated fungal strains. **E**. Representative H&E-stained colon sections of mice with DSS-induced colitis gavaged with the indicated fungal strains. **F.** A bar chart representing averages with standard deviations (SDs) of histological scores of at least five colon sections of mice gavaged with each indicated fungal strain (**p < 0.005). Control, PBS; Reference, *S. cerevisiae* ATCC 204508; Skin 1, *M. globosa* KCTC 27541; Skin 2, *M. globosa* KCTC 27777; Gut 1, *M. globosa* KCTC 37188; and Gut 2, *M. globosa* KCTC 37189.

Mice inoculated with two independent gut isolates showed a significantly reduced survival rate; nine out of 11 mice were dead on day 7 postinfection. In contrast, most mice inoculated with two independent skin isolates survived; only two out of 11 mice were dead (Fig. 4B). The negative control and the reference strain *S. cerevisiae* ATCC 204508 showed similar results to the skin isolates, confirming that only the gut isolates and not the skin isolates influenced the survival of the mice.

In addition to the survival assay, we conducted separate experiments to determine the disease activity index (DAI) of colitis, as described previously ^45^. Notably, we reduced the number of oral gavage of fungal cells to mice with DSS-induced colitis from three to two in order to maintain the survival of the host during the experiments. We found that the DAI score of the mice gavaged with the negative control, the *S. cerevisiae* reference strain, and the *M. globosa* skin isolates quickly reduced in 2–3 days after oral gavage of the fungal cells. In contrast, the DAI scores of the mice inoculated with the *M. globosa* gut isolates were relatively higher than those of the other groups, being most significant on day 2; these results indicate that DSS-induced colitis was significantly exacerbated by the gut isolates (Fig. 4C). In accordance with the results of DAI scoring, the colon length was significantly lower in the mice gavaged with the *M. globosa* gut isolates than in the other groups (Fig. 4D). Moreover, histological scoring revealed that the symptoms of colitis were greater in the mice gavaged with the *M. globosa* gut isolates than in the other groups (Fig. 4E and F).

Maintaining a proper balance of inflammatory cytokines is crucial for a healthy gut homeostasis, and the disruption of this balance can trigger the progression of diseases, including UC. During the activation of inflammatory cascades in the gut mucosa of patients with UC, the levels of various inflammatory cytokines are inappropriately elevated ^46,47^. Therefore, we measured the levels of proinflammatory cytokines, including TNF-α, IL-6, IL-12p40, IL-1β, and IL-18, in the mice gavaged with the *M. globosa* gut isolates. The results of enzyme-linked immunosorbent assay (ELISA) indicated that the cytokine levels were significantly higher in the mice inoculated with the *M. globosa* gut isolates than in those inoculated with the *M. globosa* skin isolates and the *S. cerevisiae* reference strain (Fig. 5A). These results further confirmed that the *M. globosa* gut isolates and not the skin isolates could affect inflammation and exacerbate DSS-induced colitis in the mouse model.

**Fig. 5.**
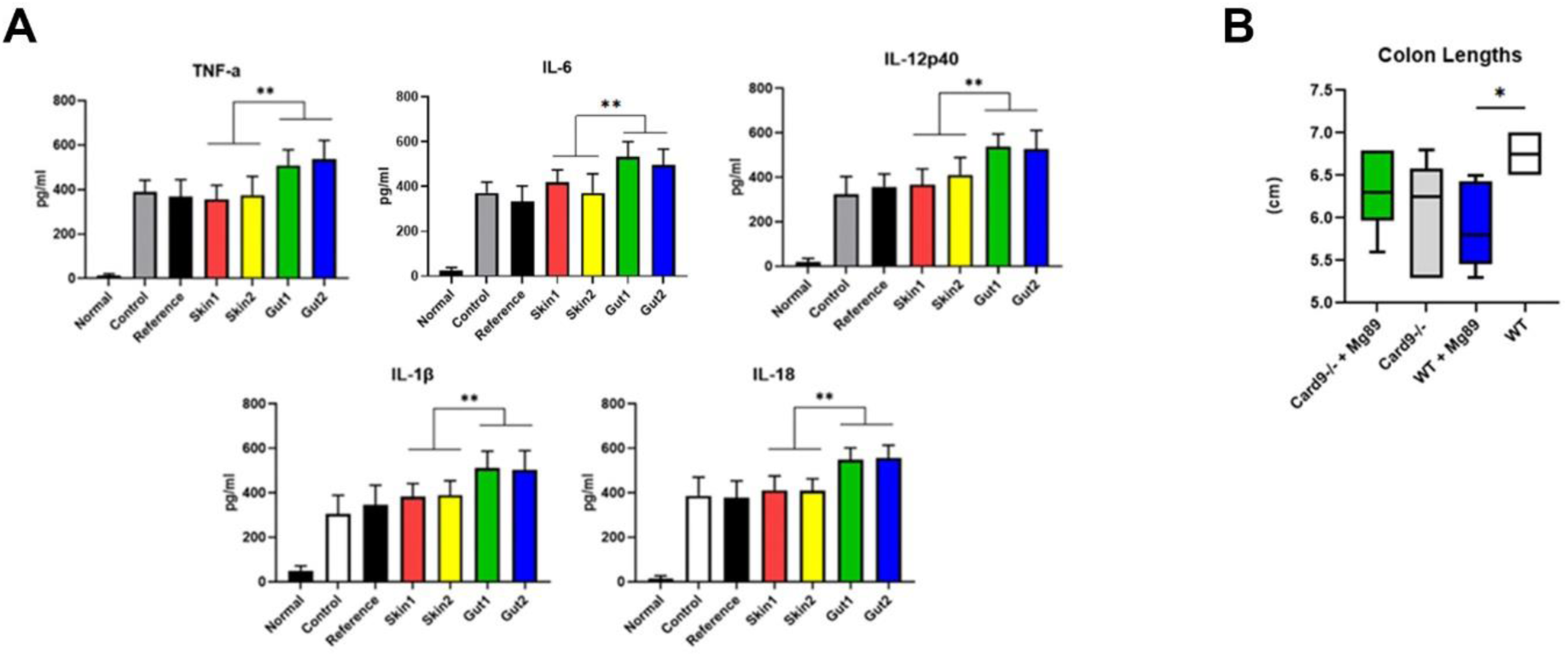
Levels of cytokines and requirement of CARD9 for *M. globosa*-associated colitis in mice. **A**. Cytokine levels were measured in mouse colonic tissue extracts. Values are averages of five independent colonic tissue extracts from mice gavaged with each indicated fungal strain. Error bars indicate SDs (**p < 0.005). Reference, *S. cerevisiae* ATCC 204508; Skin 1, *M. globosa* KCTC 27541; Skin 2, *M. globosa* KCTC 27777; Gut 1, *M. globosa* KCTC 37188; and Gut 2, *M. globosa* KCTC 37189. **B.** Comparison of colon length of Card9^−/−^ and wild-type mice gavaged with the representative *M. globosa* gut isolate KCTC 37189 (Mg89). A box plot represents the colon length of mice gavaged with either PBS or the indicated *M. globosa* strain. In each group, three mice were tested.

As mentioned earlier, Limon et al. suggested the requirement of functional CARD9 for *Malassezia-* associated colitis in a mouse model ^25^. Therefore, we assessed whether CARD9 is required for the *M. globosa* gut isolates to exacerbate DSS-induced colitis in a mouse model. The results of our experiment revealed no significant difference between Card9−/− mice gavaged with the PBD control and the *M. globosa* gut isolates; however, the wild-type mice gavaged with the same fungal strain showed significantly shortened colon length (Fig. 5B). This result confirmed the requirement of CARD9 for *M. globosa*-associated colitis in mice. Overall, our data revealed distinct pathological characteristics between the *M. globosa* gut and skin isolates and suggested that the same *Malassezia* species isolated from different host niches could play a distinct role in the host body.

## Discussion

In general, unlike bacterial diversities, fungal diversities in the gut of healthy individuals and patients with IBD exhibit significant individual variations. Moreover, different conclusions have been reported in different studies, probably because of the methodologies used and the regions and diets of the patient cohorts ^48,49^. Nevertheless, a few common features of the gut mycobiota in patients with IBD, such as an increased overall fungal load and an increased Basidiomycota/Ascomycota ratio, have been reported ^50,51^. Furthermore, several studies have reported a distinct *Malassezia* population in the gut of patients with IBD, indicating a possible role of the fungus in IBD ^21,25,50–53^. However, in our study, no significant difference was noted in the fungal diversity between the gut mucosa of patients with UC and healthy individuals. Similarly, a recent study using fecal samples of a Japanese cohort revealed no significant differences in the fungal diversity between patients with UC and healthy controls ^53^, implying that the structure of the gut mycobiota in East Asian patients with UC, in particular, is similar to that in healthy individuals.

We isolated two independent live *M. globosa* strains directly from the gut mucosa of two patients with UC but failed to obtain *M. restricta* although fungal community analysis using amplicon sequence reads indicated a significant amount of *M. restricta* in the gut mucosa of patients with UC. Failure to isolate *M. restricta* directly from the gut mucosa suggests a distinct requirement of a specific nutrient and/or a culture condition for the *M. restricta* species associated with the gut mucosa in comparison with the same species isolated from the skin. This also suggests a possible intraspecies variant of the fungus. Nevertheless, our data suggested that the *M. globosa* strains directly isolated from the gut mucosa of patients with UC exacerbated DSS-induced colitis in the mouse model. The increased disease severity was supported by the findings of increased DAI and histopathological scores, and decreased gut length, and increased production of inflammatory cytokines in the mice inoculated with the *M. globosa* gut isolates.

The existence of intraspecies variants has been reported in major human pathogenic fungi, such as *C. neoformans* and *C. albicans*. Analysis of 230 clinical isolates of *C. neoformans* from patients with HIV in South Africa revealed a close association of genotypes and phenotypes, including virulence traits of the strains, with clinical outcomes ^54^. Clinical isolates of *C. albicans* also exhibited large genetic diversity, resulting in phenotypic differences, including those in host immune responsiveness and pathogenicity, between the strains ^55–57^. In addition, the niche-specific adaptation of a *C. albicans* strain was suggested. Lemberg et al. compared transcriptomes, metabolic profiles, and pathogenicity between two *C. albicans* strains, namely strain 101, which was isolated from the oral mucosa of a healthy individual, and SC5314, a well-known laboratory standard reference strain that is also known as a hypervirulent strain. *C. albicans* strain 101 exhibited low virulence and had metabolically adapted to the oral epithelium to persist as a commensal, while the highly virulent strain SC5314 induced strong inflammation ^58^. Taking these findings into account, we compared the genome sequences of *M. globosa* strains that were directly isolated from the gut mucosa of patients with UC in this study with those of *M. globosa* strains isolated from the skin in our previous study ^36^.

Our data revealed no obvious differences in genome sequences and gene contents between the gut and skin isolates. Comparisons of gene copy number variations (CNVs) revealed the enrichment of genes involved in stress response and virulence in all *M. globosa* isolates; however, no difference was noted between the strains. CNV influences the tolerance of cells to stressful environments. In genetic studies using *S. cerevisiae*, CNV has been found to affect gene expression and cause the overexpression of a corresponding gene, directly influencing the phenotypes of cells. If the benefits outweigh the cost, CNVs become fixed; the underlying mechanism is highly specific to each strain ^59^. Our genome analysis revealed that many genes showed significant CNV across strains in the gut and skin isolates. Genes showing significant CNV are likely essential players involved in the stress tolerance, survival, and virulence of *M. globosa*, especially in the host environment. The top three genes that showed significant CNV across the strains included those associated with iron transport, RNA helicase, and the bacterial enterotoxin septicolysin homolog. Iron transport plays a vital role in the stress tolerance and virulence of pathogenic fungi ^60^. We found that the CNV of the RNA helicase gene was also variable across the strains, implying that such a gene also plays a role in stress tolerance. The homologous protein Ski2 in *C. neoformans* plays a key role in conferring resistance to azole antifungal drugs and tolerance to stressful conditions ^61^. Moreover, CNV of the gene encoding the homolog of septicolysin, which may act as a bacterial cytolytic toxin in mammalian hosts ^62,63^, was observed indicating that the gene may play a role in *M. globosa* in the host niche.

Maintaining a proper gut environment is essential to prevent disease. The gut microbiota, including the mycobiota, plays a critical role in gut homeostasis. Oxygen is a microenvironmental factor that contributes to homeostasis in the gut ecosystem and influences the metabolism and immunity of the gut and the colonization of pathogens. Several studies have demonstrated that the disruption of the oxygen gradient significantly contributes to various intestinal diseases, such as IBD and colorectal cancer ^40,41^. The existing oxygen gradient has been reported to be 5%–10% at the crypt and lumen interface and approximately 2% and 0.4% in the lumen of the ascending and sigmoid colon, respectively ^40,64^. Multiple mechanisms are involved in reducing oxygen concentrations in the gut. These include microbially derived short-chain fatty acids, such as butylate, which can promote mitochondrial β- oxidation and oxidative phosphorylation in colonocytes to increase oxygen consumption and to induce a hypoxic environment in the gut, thereby suppressing the colonization of pathogenic bacteria, such as *E. coli* and *Salmonella*. In contrast, antibiotic treatment increases oxygen concentrations, reprograms colonocytes to a state of anaerobic glycolysis from oxidative metabolism, and triggers an increase in the population of facultative pathogenic bacteria, such as *Salmonella* ^65,66^.

Compared with the hypoxic condition in the gut, normoxia is observed in the skin, especially the outmost surface of the skin where *Malassezia* dominantly resides. Therefore, we hypothesized that the *M. globosa* gut and skin isolates have distinctly adapted to the environments with different oxygen concentrations and display different physiological responsiveness to low and high concentrations of oxygen. For example, the *M. globosa* gut isolates may be more sensitive to normoxia than the skin isolates. To confirm this hypothesis, we analyzed and compared the transcriptomes of the *M. globosa* gut and skin isolates that were grown in the presence of 2% or 20% oxygen. A comparison of growth rates between the *M. globosa* gut and skin isolates under different oxygen concentrations was not possible because fungal cells form severe clumps in a liquid medium. Transcriptome analysis revealed that the genes involved in peroxisome functions and stress responses were upregulated in the *M. globosa* gut isolates in the presence of 20% oxygen, indicating that the fungal cells were highly responsive to oxygen concentrations and experienced oxidative stress under the 20% oxygen condition in comparison with the skin isolates. The enrichment of genes involved in ribosomal functions in the *M. globosa* gut isolates in the presence of 2% oxygen further supported our finding that the gut isolates may prefer a low-oxygen environment, which may resemble the environment at the gut mucosal surface.

The different transcriptome profiles noted between the gut and skin isolates may reflect the phenotypes of the strains. The *M. globosa* gut isolates showed distinct virulence levels compared with the skin isolates. The gut isolates exacerbated DSS-induced colitis in the mouse model, while the skin isolates showed similar outcomes to the negative control and the reference strain *S. cerevisiae*. Thus, although they are the same species, the *M. globosa* gut and skin isolates may have adapted to the different environment niches in the host body and may have thereby gained unique pathological characteristics.

Intraspecies diversity within the same fungal species has been commonly reported in recent studies ^55,67–69^. For example, *S. cerevisiae* and *C. albicans* isolated from the feces of patients with CD displayed different in vitro and in vivo immune responses ^68^. Moreover, *C. albicans* strains isolated from patients with irritable bowel syndrome (IBS) displayed phenotypic diversities, and genotypic analysis revealed that the strains isolated from the gut of hypersensitive IBS patients partially clustered together ^69^. However, whether the colonization of a specific body site is associated with genotypic and phenotypic diversities, including virulence, remains unclear. A recent study using 910 commensal isolates from 35 healthy donors reported extensive phenotypic diversity that was not dependent on the isolation site or participant ^57^. Furthermore, high rates of heterozygosity, structural mutation, and chromosomal variation in *C. albicans* have been reported in other studies ^70–72^. Unlike *C. albicans*, we found that the genotypes and phenotypes of *M. globosa* clinical isolates were associated with the body site. However, we used a limited number of fungal isolates. Moreover, to date, no information has been provided on the degree of heterozygosity, structural mutation, and chromosomal variation using a reasonable number of *M. globosa* clinical isolates. Therefore, further research should be conducted to analyze and confirm the relationship between a specific body site and genotypic and phenotypic variations in *Malassezia* species using a decent number of clinical isolates.

This is the first study to isolate a live *M. globosa* strain from the mucosal surface of patients with UC. We compared the genomes, transcriptomes, and virulence of the *M. globosa* gut isolates with those of the skin isolates and noted distinct transcriptome profiles between the gut and skin isolates, suggesting that the gut isolates are more susceptible to normoxia (approximately 20% oxygen) than the skin isolates. These results imply that the *M. globosa* gut isolates have evolutionally adapted to hypoxia, which reflects the environment in the gut mucosa. Using the DSS-induced colitis mouse model, we found that the *M. globosa* gut isolates significantly exacerbated colitis. However, no difference was noted between the skin isolates and the negative control. These findings further confirmed the presence of different phenotypes between the gut and skin isolates and suggested that *M. globosa* plays a role in the gut mucosa in the mouse model. Overall, our data provide novel insights into the pivotal role of *M. globosa* in the pathogenesis of IBD, highlighting the potential influence of niche-specific adaptations on the virulence of this fungus. The findings also provide crucial information on the complex interplay between the members of the gut mycobiota and host health.

## Methods

### Fungal strains

*S. cerevisiae* ATCC 204508 was used as a negative reference strain and grown overnight at 30°C in yeast extract peptone dextrose (YPD) medium. *M. globosa* KCTC 27541 and KCTC 27777, which were isolated from the skin lesions on the scalp of patients with seborrheic dermatitis in our previous study ^73^, were used as skin isolates and compared with the gut isolates throughout the study. Two independent *M. globosa* strains were isolated from the gut mucosa of two different patients with UC during colonoscopy, as described below. The strains were identified by polymerase chain reaction (PCR) using the primers ITS4 (TCCTCCGCTTATTGATATGC) and ITS5 (GGAAGTAAAAGTCGTAACAAGG) and by sequencing. They were deposited at the KCTC and designated *M. globosa* KCTC 37188 and KCTC 37189, respectively ^74^. *M. globosa* strains were cultured at 34°C on LNA medium for three days and used throughout the study.

### Collection of mucosal lavage samples

This study was conducted in adult patients with UC who were aged 19 years and older and attended Chung-Ang University Hospital between October 2021 and August 2022 and healthy controls. Samples from patients with UC were obtained during colonoscopic examinations for clinical reasons. In total, 20 mL of distilled water was injected twice or thrice into the colonic mucosa through the instrument channel of a colonoscope, followed by collection (approximately 50 mL) through the suction channel. These mucosal lavage samples were collected separately from areas with and without inflammation from the same patient. Patients’ clinical data, including demographic factors, disease characteristics, endoscopic findings at the time of sample collection, and sampling sites, were recorded. The surgical history of bowel resection and medication history, including the use of steroids, immunomodulators, and biologics, were also recorded. Healthy controls were individuals who visited the gastroenterology clinic to undergo colonoscopy for routine check-ups and had not been diagnosed with CD, UC, or functional gastrointestinal disorders based on the ROME IV criteria ^75^. Participants with serious chronic conditions that could potentially impact the results of this study, such as uncontrolled hypertension, diabetes, pulmonary tuberculosis, liver cirrhosis, and chronic kidney disease, as well as those who had received systemic or local antifungal agents within the past month were excluded. Mucosal lavage samples were obtained from healthy controls using the same procedure as that used for patients with UC; the samples were obtained from a randomly selected area of the colon during routine colonoscopic examinations. This study was approved by the Institutional Review Board of Chung-Ang University Hospital (IRB No. 2002-024-441). Written informed consent was obtained from each subject before participation in the study.

### DNA extraction and fungal isolation

The mucosal lavage samples were centrifuged at 6,000 rpm for 30 min at room temperature, and the pellet was suspended in 1 mL of phosphate-buffered saline (PBS). DNA was extracted from 500 μL of the suspension, as described previously ^76^, and used for fungal community analysis. The remaining suspension was spread on LNA medium containing antibiotics (gentamicin 100 μg/mL, chloramphenicol 50 μg/mL) to restrict the growth of bacterial cells and cultured for 2 weeks at 34°C in the presence of 2% oxygen. A single colony was obtained and subjected to species identification by PCR amplification using ITS4 and ITS5 primers and by sequencing, as described above.

### Library construction, sequencing, and data analysis of the mycobiome

The sequencing libraries were prepared according to the Illumina Metagenomic Sequencing Library protocols to amplify the ITS3 and ITS4 regions. The input gDNA (10 ng) was PCR amplified using 5× reaction buffer, 1 mM of dNTP mix, 500 nM each of the universal F/R PCR primers, and the Herculase II fusion DNA polymerase (Agilent Technologies, Santa Clara, CA, USA). The cycle condition for the first PCR was 3 min at 95°C for heat activation; 35 cycles of 30 s at 95°C, 30 s at 55°C, and 30 s at 72°C; and a 5-min final extension at 72°C. The universal primer pair with the Illumina adapter overhang sequences used for the first amplification was as follows:

ITS3 amplicon PCR forward primer:

5′-TCGTCGGCAGCGTCAGATGTGTATAAGAGACAGGCATCGATGAAGAACGCAGC-3′

ITS4 amplicon PCR reverse primer:

5′-GTCTCGTGGGCTCGGAGATGTGTATAAGAGACAGTCCTCCGCTTATTGATATGC-3′

The first PCR product was purified using AMPure beads (Agencourt Bioscience, Beverly, MA, USA). Following purification, 10 µL of the first PCR product was PCR amplified to construct the final library containing an index using the Nextera XT Indexed Primer. The cycle condition for the second PCR was the same as that for the first PCR, except for 10 cycles. The PCR product was purified using AMPure beads. The final purified product was then quantified using qPCR, according to the qPCR Quantification Protocol Guide (KAPA Library Quantification Kit for Illumina Sequencing platforms), and qualified using the TapeStation D1000 ScreenTape (Agilent Technologies, Waldbronn, Germany). The product was then sequenced using the MiSeq™ platform (Illumina, San Diego, USA).

Generated raw reads were trimmed based on the quality, and adapters were removed using Trimmomatic v.0.36 ^77^. The cleaned paired-end reads were merged using PEAR v.0.9.6 ^78^, and the results were imported to the QIIME 2 pipeline v.2020.8 ^79^. Taxonomic assignment was performed using the Targeted Host-associated Fungi (THF) v1.6.1 mycobiome database ^80^. α-diversity and β-diversity metrics were measured in QIIME 2 using the default parameters.

### Genome sequencing and analysis

Genomic DNA of the *M. globosa* gut isolates (KCTC 37188 and KCTC 37189) and skin isolates (KCTC 27541 and KCTC 27777) was extracted using the Wizard HMW DNA Extraction Kit (Promega, Madison, WI, USA), according to the manufacturer’s recommendations. The sequencing library was constructed using the TruSeq DNA PCR-Free Library Prep Kit (Illumina, San Diego, CA, USA), according to the manufacturer’s instructions. NGS was performed using NovaSeq 6000 (Illumina, San Diego, CA, USA) on a 151-bp paired-end platform.

The generated raw data were quality filtered and adapter trimmed by Trimmomatic v.0.36 ^77^ using default parameters. Cleaned reads were assembled using the CLC Genomics Workbench v20 (Qiagen, Germantown, MD, USA). From the resulting contigs, gene prediction and annotation were performed by FunGap v.1.1.0 ^81^ using the RNA-Seq data of each strain. Orthologs were grouped by DIAMOND v.2.0.6 ^82^ with the parameter “-id 50” and MCL 14–137 with the parameter “-I 1.5” ^83^. ANI values were analyzed using the ANIb method based on BLAST via JSpecies-WS ^84^.

### RNA extraction and transcriptome analysis

*M. globosa* strains were cultured in LNA medium for 3 days at 34°C in the presence of 2% or 20% oxygen (CO_2_/O_2_ incubator, Vision Scientific, Daejun, Korea). RNA extraction was performed using TRIzol™ Reagent (Invitrogen, Carlsbad, CA, USA), according to the manufacturer’s instructions. The extracted RNA was further treated using the TURBO DNA-Free™ Kit (Ambion, Austin, TX, USA), according to the manufacturer’s instructions. Sequencing libraries for transcriptome analysis (RNA-Seq) were constructed using the TruSeq Stranded mRNA Library Prep Kit (Illumina, San Diego, CA, USA), according to the manufacturer’s instructions. Sequencing was performed using NovaSeq 6000 (Illumina, San Diego, CA, USA), according to the manufacturer’s instructions, and 101-bp paired-end reads were generated. Raw data were trimmed using the same procedure as that used for genome sequencing. Cleaned reads were mapped to the assembled genome using hisat2 v.2.2.1 ^85^. The reads that were mapped to each CDS were counted using featureCounts ^86^. Finally, the counts from each CDS were normalized and statistically analyzed using the DeSeq2 package ^87^. Functional enrichment analysis against differentially expressed genes under each condition was performed using DAVID Bioinformatics Resources ^88^.

### Determination of survival rate and DAI using a DSS-induced colitis mouse model

*M. globosa* gut isolates (KCTC 37188 and KCTC 37189) and skin isolates (KCTC 27541 and KCTC 27777) were cultured on LNA medium for three days at 34°C. *S. cerevisiae* ATCC 204508 was cultured on YPD agar for two days at 30℃. The fungal cells were scraped from the agar plate, washed in PBS, and filtered using a 40-µm-diameter filter. The filtered cells were then counted and resuspended to a concentration of 1 × 10^8^ fungal cells/mL. Eleven female C57BL/6 mice (Koatech, Pyeongtaek, Korea) per group were used.

A DSS-induced colitis mouse model was developed using 6-week-old female C57BL/6 mice (Koatech, Pyeongtaek, Korea), as described previously ^89^. Colitis was induced using 2.5% DSS (molecular weight: 36,000–50,000 kDa, MP Biomedicals, Santa Ana, CA, USA) for five days, followed by water for two days (i.e., a total of seven days). On day eight, 100 μL of PBS containing 1 × 10^7^ fungal cells was orally gavaged to the colitis-induced mice every other day for five days (three injections in total), and the survival of the mice was monitored daily. Five mice were sacrificed on day four after two injections, and the colon length and DAI score were determined as reported previously ^45^. During the experiment, the stool and weight of each mouse were assessed and scored daily. All procedures for animal experiments were reviewed and approved by the Institutional Animal Care and Use Committee of Hanyang University (protocol 2022-0103). All animal experiments followed the guidelines provided by the Korean Food and Drug Administration.

### ELISA

The levels of TNF-α, IL-6, IL-12p40, IL-1β, and IL-18 were measured in mouse colonic tissue extracts by using the BD OptEIA ELISA System (BD Biosciences, San Jose, CA, USA). All assays were performed as per the manufacturer’s instructions.

### Histological assessment

For immunohistochemical assessment of tissue sections, the distal colonic tissues of mice were fixed in 10% formalin and embedded in paraffin. The paraffin-embedded sections (4 μm thick) were sliced and stained with hematoxylin and eosin (H&E). Histopathological scores and H-scores (a semiquantitative measure) were assessed ^90^. For this purpose, a board-certified pathologist (Dr. Min-Kyung Kim, Kim Min-Kyung Pathology Clinic, Seoul, Korea) independently evaluated each organ section without prior knowledge of the treatment groups.

### Determination of colon length of DSS-induced CARD9-knockout mice

Female wild-type C57BL/6J mice were purchased from Jackson Laboratories. Wild-type and CARD9-knockout mice were cohoused for 2 weeks before initiating the experiment. The mice were housed under specific pathogen-free conditions at the Cedars Sinai Medical Center animal facility and handled in accordance with approved IACUC standards and procedures. The mice were given 3.5% DSS (MP Biomedicals) for seven days and then given water for by three days (i.e., 10 days in total). On day 11, 100 µL of PBS containing 1 × 10^7^ fungal cells was orally gavaged into each mouse every other day for four days (two injections in total), and the colon length was measured.

### Statistical analysis

Statistical analysis was performed using one-way ANOVA with Tukey’s multiple comparison test on GraphPad Prism version 9.5.1. Statistical significance was defined as a p value of ≤0.05.

## Data accessibility

The ITS2 sequence reads, genome sequences, and RNA sequencing data are available in the NCBI SRA under the BioProject accession number PRJNA1013764.

## Author Contributions

## Supplementary Materials

**Table S1.** List of patients (healthy controls and patients with UC): age, sex, disease information, and treatment data.

**Table S2.** Live fungal strains obtained in this study.

**Fig. S1.** α- and β-diversity analyses.

## References

1 Maloy, K. J. & Powrie, F. Intestinal homeostasis and its breakdown in inflammatory bowel disease. Nature 474, 298–306, doi:10.1038/nature10208 (2011).

2 Ramos, G. P. & Papadakis, K. A. Mechanisms of Disease: Inflammatory Bowel Diseases. Mayo Clin Proc 94, 155–165, doi:10.1016/j.mayocp.2018.09.013 (2019).

3 Neurath, M. F. Targeting immune cell circuits and trafficking in inflammatory bowel disease. Nat Immunol 20, 970–979, doi:10.1038/s41590-019-0415-0 (2019).

4 Buie, M. J. et al. Global hospitalization trends for Crohn’s disease and ulcerative colitis in the 21st Century: A Systematic Review With Temporal Analyses. Clin Gastroenterol Hepatol 21, 2211–2221, doi:10.1016/j.cgh.2022.06.030 (2023).

5 Ashton, J. J. & Beattie, R. M. Inflammatory bowel disease: recent developments. Arch Dis Child, doi:10.1136/archdischild-2023-325668 (2023).

6 Lee, J. W. & Eun, C. S. Inflammatory bowel disease in Korea: epidemiology and pathophysiology. Korean J Intern Med 37, 885–894, doi:10.3904/kjim.2022.138 (2022).

7 Kim, D. H. & Cheon, J. H. Pathogenesis of Inflammatory Bowel Disease and Recent Advances in Biologic Therapies. Immune Netw 17, 25–40, doi:10.4110/in.2017.17.1.25 (2017).

8 Zhao, Q. & Maynard, C. L. Mucus, commensals, and the immune system. Gut Microbes 14, 2041342, doi:10.1080/19490976.2022.2041342 (2022).

9 Shen, Z. H. et al. Relationship between intestinal microbiota and ulcerative colitis: Mechanisms and clinical application of probiotics and fecal microbiota transplantation. World J Gastroenterol 24, 5–14, doi:10.3748/wjg.v24.i1.5 (2018).

10 Andoh, A. & Nishida, A. Alteration of the Gut Microbiome in Inflammatory Bowel Disease. Digestion 104, 16–23, doi:10.1159/000525925 (2023).

11 Benech, N. & Sokol, H. Targeting the gut microbiota in inflammatory bowel diseases: where are we? Curr Opin Microbiol 74, 102319, doi:10.1016/j.mib.2023.102319 (2023).

12 Underhill, D. M. & Braun, J. Fungal microbiome in inflammatory bowel disease: a critical assessment. J Clin Invest 132, e155786, doi:10.1172/JCI155786 (2022).

13 Iliev, I. D. et al. Interactions between commensal fungi and the C-type lectin receptor Dectin-1 influence colitis. Science 336, 1314–1317, doi:10.1126/science.1221789 (2012).

14 Leonardi, I. et al. Fungal Trans-kingdom Dynamics Linked to Responsiveness to Fecal Microbiota Transplantation (FMT) Therapy in Ulcerative Colitis. Cell Host Microbe 27, 823–829 e823, doi:10.1016/j.chom.2020.03.006 (2020).

15 Ianiri, G., LeibundGut-Landmann, S. & Dawson, T. L., Jr. Malassezia: A Commensal, Pathogen, and Mutualist of Human and Animal Skin. Annu Rev Microbiol 76, 757–782, doi:10.1146/annurev-micro-040820-010114 (2022).

16 Brayer, K. J. et al. The immune response to a fungus in pancreatic cancer samples. bioRxiv, doi:10.1101/2023.03.28.534606 (2023).

17 Alonso, R., Pisa, D., Aguado, B. & Carrasco, L. Identification of Fungal Species in Brain Tissue from Alzheimer’s Disease by Next-Generation Sequencing. J Alzheimers Dis 58, 55–67, doi:10.3233/JAD-170058 (2017).

18 Alonso, R., Pisa, D., Fernandez-Fernandez, A. M. & Carrasco, L. Infection of Fungi and Bacteria in Brain Tissue From Elderly Persons and Patients With Alzheimer’s Disease. Front Aging Neurosci 10, 159, doi:10.3389/fnagi.2018.00159 (2018).

19 Oh, J. et al. Biogeography and individuality shape function in the human skin metagenome. Nature 514, 59–64, doi:10.1038/nature13786 (2014).

20 Shuai, M. et al. Mapping the human gut mycobiome in middle-aged and elderly adults: multiomics insights and implications for host metabolic health. Gut 71, 1812–1820, doi:10.1136/gutjnl-2021-326298 (2022).

21 Liguori, G. et al. Fungal Dysbiosis in Mucosa-associated Microbiota of Crohn’s Disease Patients. J Crohns Colitis 10, 296–305, doi:10.1093/ecco-jcc/jjv209 (2016).

22 Maas, E., Penders, J. & Venema, K. Fungal-Bacterial Interactions in the Human Gut of Healthy Individuals. J Fungi (Basel) 9, doi:10.3390/jof9020139 (2023).

23 Nagata, R. et al. Transmission of the major skin microbiota, Malassezia, from mother to neonate. Pediatr Int 54, 350–355, doi:10.1111/j.1442-200X.2012.03563.x (2012).

24 Wang, Y. R. et al. Infant Mode of Delivery Shapes the Skin Mycobiome of Prepubescent Children. Microbiol Spectr 10, e0226722, doi:10.1128/spectrum.02267-22 (2022).

25 Limon, J. J. et al. Malassezia is associated with Crohn’s disease and exacerbates colitis in mouse models. Cell Host Microbe 25, 377–388 e376, doi:10.1016/j.chom.2019.01.007 (2019).

26 Hartjes, L. & Ruland, J. CARD9 Signaling in Intestinal Immune Homeostasis and Oncogenesis. Front Immunol 10, 419, doi:10.3389/fimmu.2019.00419 (2019).

27 Gueho, E., Midgley, G. & Guillot, J. The genus Malassezia with description of four new species. Antonie Van Leeuwenhoek 69, 337–355, doi:10.1007/BF00399623 (1996).

28 Zhu, B. et al. Multi-omics analysis of niche specificity provides new insights into ecological adaptation in bacteria. ISME J 10, 2072–2075, doi:10.1038/ismej.2015.251 (2016).

29 Kido, Y. et al. Niche-specific adaptation of Lactobacillus helveticus strains isolated from malt whisky and dairy fermentations. Microb Genom 7, 000560, doi:10.1099/mgen.0.000560 (2021).

30 Dettman, J. R. & Kassen, R. Evolutionary Genomics of Niche-Specific Adaptation to the Cystic Fibrosis Lung in Pseudomonas aeruginosa. Mol Biol Evol 38, 663–675, doi:10.1093/molbev/msaa226 (2021).

31 Fillinger, R. J. & Anderson, M. Z. Seasons of change: Mechanisms of genome evolution in human fungal pathogens. Infect Genet Evol 70, 165–174, doi:10.1016/j.meegid.2019.02.031 (2019).

32 Alves, R., et al. Adapting to survive: How *Candida* overcomes host-imposed constraints during human colonization. Plos Pathogens 16, doi:ARTN e1008478 10.1371/journal.ppat.1008478 (2020).

33 Zmora, N. et al. Personalized Gut Mucosal Colonization Resistance to Empiric Probiotics Is Associated with Unique Host and Microbiome Features. Cell 174, 1388–1405 e1321, doi:10.1016/j.cell.2018.08.041 (2018).

34 Worsley, S. F. et al. Assessing the causes and consequences of gut mycobiome variation in a wild population of the Seychelles warbler. Microbiome 10, 242, doi:10.1186/s40168-022-01432-7 (2022).

35 Hoffmann, C. et al. Archaea and fungi of the human gut microbiome: correlations with diet and bacterial residents. PLoS One 8, e66019, doi:10.1371/journal.pone.0066019 (2013).

36 Lee, Y. W., Lee, S. Y., Lee, Y. & Jung, W. H. Evaluation of Expression of Lipases and Phospholipases of Malassezia restricta in Patients with Seborrheic Dermatitis. Ann Dermatol 25, 310–314, doi:10.5021/ad.2013.25.3.310 (2013).

37 Cho, Y. J. et al. Genome of Malassezia arunalokei and Its Distribution on Facial Skin. Microbiol Spectr 10, e0050622, doi:10.1128/spectrum.00506-22 (2022).

38 Wu, G. et al. Genus-Wide Comparative Genomics of Malassezia Delineates Its Phylogeny, Physiology, and Niche Adaptation on Human Skin. PLoS Genet 11, e1005614, doi:10.1371/journal.pgen.1005614 (2015).

39 Ianiri, G. et al. HGT in the human and skin commensal Malassezia: A bacterially derived flavohemoglobin is required for NO resistance and host interaction. Proc Natl Acad Sci U S A 117, 15884–15894, doi:10.1073/pnas.2003473117 (2020).

40 Singhal, R. & Shah, Y. M. Oxygen battle in the gut: Hypoxia and hypoxia-inducible factors in metabolic and inflammatory responses in the intestine. J Biol Chem 295, 10493–10505, doi:10.1074/jbc.REV120.011188 (2020).

41 Ramakrishnan, S. K. & Shah, Y. M. Role of Intestinal HIF-2alpha in Health and Disease. Annu Rev Physiol 78, 301–325, doi:10.1146/annurev-physiol-021115-105202 (2016).

42 Lin, N. X., He, R. Z., Xu, Y. & Yu, X. W. Augmented peroxisomal ROS buffering capacity renders oxidative and thermal stress cross-tolerance in yeast. Microb Cell Fact 20, 131, doi:10.1186/s12934-021-01623-1 (2021).

43 Schrader, M. & Fahimi, H. D. Peroxisomes and oxidative stress. Biochim Biophys Acta 1763, 1755-1766, doi:10.1016/j.bbamcr.2006.09.006 (2006).

44 Hsu, J. W. et al. Unfolded protein response regulates yeast small GTPase Arl1p activation at late Golgi via phosphorylation of Arf GEF Syt1p. Proc Natl Acad Sci U S A 113, E1683–1690, doi:10.1073/pnas.1518260113 (2016).

45 Cooper, H. S., Murthy, S. N., Shah, R. S. & Sedergran, D. J. Clinicopathologic study of dextran sulfate sodium experimental murine colitis. Lab Invest 69, 238–249 (1993).

46 Muzes, G., Molnar, B., Tulassay, Z. & Sipos, F. Changes of the cytokine profile in inflammatory bowel diseases. World J Gastroenterol 18, 5848–5861, doi:10.3748/wjg.v18.i41.5848 (2012).

47 Friedrich, M., Pohin, M. & Powrie, F. Cytokine Networks in the Pathophysiology of Inflammatory Bowel Disease. Immunity 50, 992–1006, doi:10.1016/j.immuni.2019.03.017 (2019).

48 Ost, K. S. & Round, J. L. Commensal fungi in intestinal health and disease. Nat Rev Gastroenterol Hepatol, doi:10.1038/s41575-023-00816-w (2023).

49 Nash, A. K. et al. The gut mycobiome of the Human Microbiome Project healthy cohort. Microbiome 5, 153, doi:10.1186/s40168-017-0373-4 (2017).

50 Sokol, H. et al. Fungal microbiota dysbiosis in IBD. Gut 66, 1039–1048, doi:10.1136/gutjnl-2015-310746 (2017).

51 Acar, C. et al. Composition of the colon microbiota in the individuals with inflammatory bowel disease and colon cancer. Folia Microbiol (Praha), doi:10.1007/s12223-023-01072-w (2023).

52 Cimicka, J., Riegert, J., Kavkova, M. & Cerna, K. Intestinal mycobiome associated with diagnosis of inflammatory bowel disease based on tissue biopsies. Med Mycol 60, myab076, doi:10.1093/mmy/myab076 (2022).

53 Imai, T. et al. Characterization of fungal dysbiosis in Japanese patients with inflammatory bowel disease. J Gastroenterol 54, 149–159, doi:10.1007/s00535-018-1530-7 (2019).

54 Beale, M. A. et al. Genotypic Diversity Is Associated with Clinical Outcome and Phenotype in Cryptococcal Meningitis across Southern Africa. PLoS Negl Trop Dis 9, e0003847, doi:10.1371/journal.pntd.0003847 (2015).

55 Marakalala, M. J. et al. Differential adaptation of Candida albicans in vivo modulates immune recognition by dectin-1. PLoS Pathog 9, e1003315, doi:10.1371/journal.ppat.1003315 (2013).

56 Hirakawa, M. P. et al. Genetic and phenotypic intra-species variation in Candida albicans. Genome Res 25, 413–425, doi:10.1101/gr.174623.114 (2015).

57 Anderson, F. M. et al. Candida albicans selection for human commensalism results in substantial within-host diversity without decreasing fitness for invasive disease. PLoS Biol 21, e3001822, doi:10.1371/journal.pbio.3001822 (2023).

58 Lemberg, C. et al. Candida albicans commensalism in the oral mucosa is favoured by limited virulence and metabolic adaptation. PLoS Pathog 18, e1010012, doi:10.1371/journal.ppat.1010012 (2022).

59 Robinson, D. et al. Gene-by-environment interactions influence the fitness cost of gene copy-number variation in yeast. G3 (Bethesda), doi:10.1093/g3journal/jkad159 (2023).

60 Bairwa, G., Hee Jung, W. & Kronstad, J. W. Iron acquisition in fungal pathogens of humans. Metallomics 9, 215–227, doi:10.1039/c6mt00301j (2017).

61 Li, C. et al. The RNA helicase Ski2 in the fungal pathogen Cryptococcus neoformans highlights key roles in azoles resistance and stress tolerance. Med Mycol 60, myac083, doi:10.1093/mmy/myac083 (2022).

62 Billington, S. J., Jost, B. H. & Songer, J. G. Thiol-activated cytolysins: structure, function and role in pathogenesis. FEMS Microbiol Lett 182, 197–205, doi:10.1016/s0378-1097(99)00536-4 (2000).

63 Acosta, J. et al. Multidrug-resistant Acinetobacter baumannii harboring OXA-24 carbapenemase, Spain. Emerg Infect Dis 17, 1064–1067, doi:10.3201/eid/1706.091866 (2011).

64 He, G. et al. Noninvasive measurement of anatomic structure and intraluminal oxygenation in the gastrointestinal tract of living mice with spatial and spectral EPR imaging. Proc Natl Acad Sci U S A 96, 4586–4591, doi:10.1073/pnas.96.8.4586 (1999).

65 Gillis, C. C. et al. Dysbiosis-Associated Change in Host Metabolism Generates Lactate to Support Salmonella Growth. Cell Host Microbe 23, 570, doi:10.1016/j.chom.2018.03.013 (2018).

66 Pral, L. P., Fachi, J. L., Correa, R. O., Colonna, M. & Vinolo, M. A. R. Hypoxia and HIF-1 as key regulators of gut microbiota and host interactions. Trends Immunol 42, 604–621, doi:10.1016/j.it.2021.05.004 (2021).

67 Cavalieri, D. et al. Genomic and Phenotypic Variation in Morphogenetic Networks of Two Candida albicans Isolates Subtends Their Different Pathogenic Potential. Front Immunol 8, 1997, doi:10.3389/fimmu.2017.01997 (2017).

68 Di Paola, M. et al. Comparative immunophenotyping of Saccharomyces cerevisiae and Candida spp. strains from Crohn’s disease patients and their interactions with the gut microbiome. J Transl Autoimmun 3, 100036, doi:10.1016/j.jtauto.2020.100036 (2020).

69 van Thiel, I. A. M. et al. Genetic and phenotypic diversity of fecal Candida albicans strains in irritable bowel syndrome. Sci Rep 12, 5391, doi:10.1038/s41598-022-09436-x (2022).

70 MacCallum, D. M. et al. Property differences among the four major Candida albicans strain clades. Eukaryot Cell 8, 373–387, doi:10.1128/EC.00387-08 (2009).

71 Forche, A. et al. Selection of Candida albicans trisomy during oropharyngeal infection results in a commensal-like phenotype. PLoS Genet 15, e1008137, doi:10.1371/journal.pgen.1008137 (2019).

72 Ene, I. V., Bennett, R. J. & Anderson, M. Z. Mechanisms of genome evolution in Candida albicans. Curr Opin Microbiol 52, 47–54, doi:10.1016/j.mib.2019.05.001 (2019).

73 Park, M., Jung, W. H., Han, S. H., Lee, Y. H. & Lee, Y. W. Characterisation and Expression Analysis of MrLip1, a Class 3 Family Lipase of Malassezia restricta. Mycoses 58, 671–678, doi:10.1111/myc.12412 (2015).

74 Irinyi, L. et al. International Society of Human and Animal Mycology (ISHAM)-ITS reference DNA barcoding database--the quality controlled standard tool for routine identification of human and animal pathogenic fungi. Med Mycol 53, 313–337, doi:10.1093/mmy/myv008 (2015).

75 Drossman, D. A. & Hasler, W. L. Rome IV-Functional GI Disorders: Disorders of Gut-Brain Interaction. Gastroenterology 150, 1257–1261, doi:10.1053/j.gastro.2016.03.035 (2016).

76 Dore, J., et al. IHMS_SOP 06 V1: Standard operating procedure for fecal samples DNA extraction, Protocol Q. International Human Microbiome Standards (2015).

77 Bolger, A. M., Lohse, M. & Usadel, B. Trimmomatic: a flexible trimmer for Illumina sequence data. Bioinformatics 30, 2114–2120, doi:10.1093/bioinformatics/btu170 (2014).

78 Zhang, J., Kobert, K., Flouri, T. & Stamatakis, A. PEAR: a fast and accurate Illumina Paired-End reAd mergeR. Bioinformatics 30, 614–620, doi:10.1093/bioinformatics/btt593 (2014).

79 Bolyen, E. et al. Reproducible, interactive, scalable and extensible microbiome data science using QIIME 2. Nat Biotechnol 37, 852–857, doi:10.1038/s41587-019-0209-9 (2019).

80 Tang, J., Iliev, I. D., Brown, J., Underhill, D. M. & Funari, V. A. Mycobiome: Approaches to analysis of intestinal fungi. J Immunol Methods 421, 112–121, doi:10.1016/j.jim.2015.04.004 (2015).

81 Min, B., Grigoriev, I. V. & Choi, I. G. FunGAP: Fungal Genome Annotation Pipeline using evidence-based gene model evaluation. Bioinformatics 33, 2936–2937, doi:10.1093/bioinformatics/btx353 (2017).

82 Buchfink, B., Xie, C. & Huson, D. H. Fast and sensitive protein alignment using DIAMOND. Nat Methods 12, 59–60, doi:10.1038/nmeth.3176 (2015).

83 Dongen, S. V. Graph Clustering Via a Discrete Uncoupling Process. SIAM J Matrix Anal and Appl 30, 121–141, doi:10.1137/040608635 (2008).

84 Richter, M., Rossello-Mora, R., Oliver Glockner, F. & Peplies, J. JSpeciesWS: a web server for prokaryotic species circumscription based on pairwise genome comparison. Bioinformatics 32, 929–931, doi:10.1093/bioinformatics/btv681 (2016).

85 Kim, D., Paggi, J. M., Park, C., Bennett, C. & Salzberg, S. L. Graph-based genome alignment and genotyping with HISAT2 and HISAT-genotype. Nat Biotechnol 37, 907–915, doi:10.1038/s41587-019-0201-4 (2019).

86 Liao, Y., Smyth, G. K. & Shi, W. The Subread aligner: fast, accurate and scalable read mapping by seed-and-vote. Nucleic Acids Res 41, e108, doi:10.1093/nar/gkt214 (2013).

87 Love, M. I., Huber, W. & Anders, S. Moderated estimation of fold change and dispersion for RNA-seq data with DESeq2. Genome Biol 15, 550, doi:10.1186/s13059-014-0550-8 (2014).

88 Sherman, B. T. et al. DAVID: a web server for functional enrichment analysis and functional annotation of gene lists (2021 update). Nucleic Acids Res, doi:10.1093/nar/gkac194 (2022).

89 Kim, J. S. et al. Mito-TIPTP Increases Mitochondrial Function by Repressing the Rubicon-p22phox Interaction in Colitis-Induced Mice. Antioxidants (Basel) 10, doi:10.3390/antiox10121954 (2021).

90 Kennedy, R. J. et al. Interleukin 10-deficient colitis: new similarities to human inflammatory bowel disease. Br J Surg 87, 1346–1351, doi:10.1046/j.1365-2168.2000.01615.x (2000).

