## Supplemental Figure S1, Table S1, Table S2 for "Live *Malassezia* strains isolated from the mucosa of patients with ulcerative colitis": Figure S1.pdf

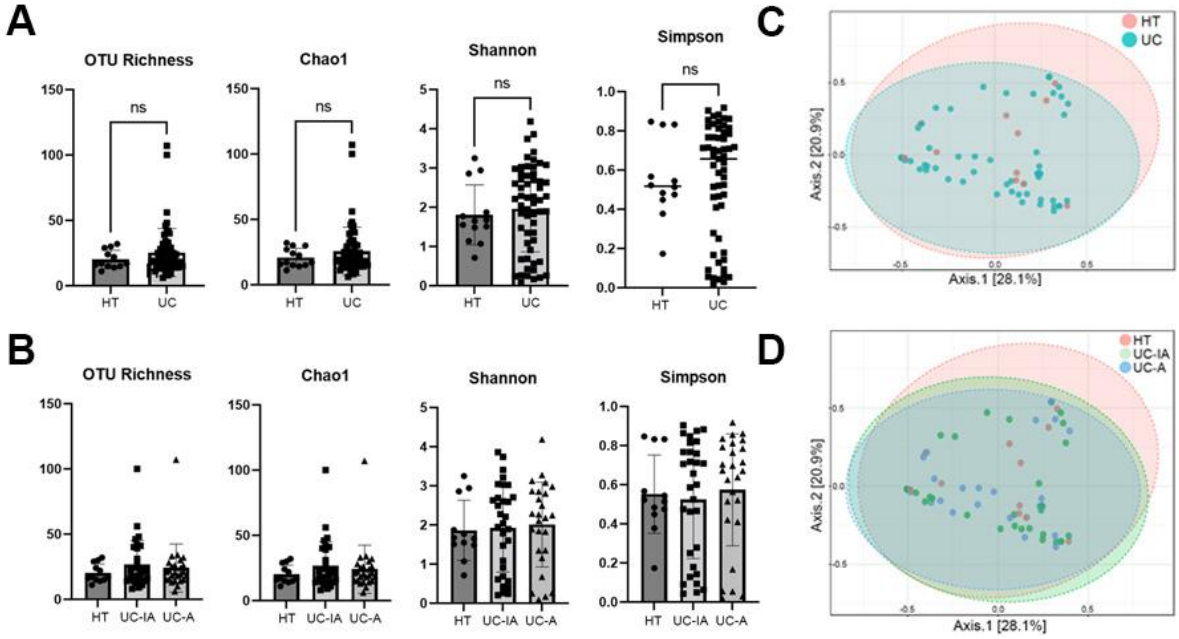


**Fig. S1.** α- and β-diversity analyses. Box plots of α-diversity distribution (richness and Chao, Shannon, and Simpson diversities). **A.** Healthy controls (HT) versus patients with UC. **B.** Healthy controls (HT) and the sites with inflammation (UC-A) versus those without inflammation (UC-IA) from patients with UC. PCoA plots of β-diversity. **C.** Healthy controls (HT) versus patients with UC. **D.** Healthy controls (HT) and the sites with inflammation (UC-A) versus those without inflammation (UC-IA) from patients with UC.
