## Supplemental Figure S1, Table S1, Table S2 for "Live *Malassezia* strains isolated from the mucosa of patients with ulcerative colitis": Table S1. List of patients.pdf

**Table S1. List of patients included in the current study.**

| Sample ID | Gender/Age | HT/PT | Sampling site | Inflammation (-/+) |
|---|---|---|---|---|
| 13 | M/40 | Ulcerative colitis | Sigmoid colon | + |
| 14 | M/40 | Ulcerative colitis | Sigmoid colon | - |
| *19 | F/62 | Ulcerative colitis | Descending colon | - |
| 20 | M/47 | Ulcerative colitis | Descending colon | - |
| 21 | M/47 | Ulcerative colitis | Sigmoid colon | - |
| *22 | F/71 | Ulcerative colitis | Descending colon | - |
| 25 | M/41 | Ulcerative colitis | Transverse colon | - |
| 26 | M/41 | Ulcerative colitis | Sigmoid colon | + |
| 27 | M/66 | Ulcerative colitis | Sigmoid colon | + |
| 28 | M/66 | Ulcerative colitis | Rectum | - |
| 31 | F/37 | Ulcerative colitis | Rectum | + |
| 32 | F/37 | Ulcerative colitis | Descending colon | - |
| 35 | F/52 | Ulcerative colitis | Sigmoid colon | + |
| 36 | F/52 | Ulcerative colitis | Descending colon | - |
| 38 | F/20 | Ulcerative colitis | Sigmoid colon | + |
| 39 | F/20 | Ulcerative colitis | Transverse colon | - |
| 42 | M/62 | Ulcerative colitis | Descending colon | + |
| 43 | M/62 | Ulcerative colitis | Sigmoid colon | - |
| 50 | M/63 | Ulcerative colitis | Sigmoid colon | + |
| 51 | M/63 | Ulcerative colitis | Transverse colon | - |
| 54 | M/28 | Ulcerative colitis | Descending colon | + |
| 55 | M/28 | Ulcerative colitis | Rectum | - |
| 69 | F/33 | Ulcerative colitis | Transverse colon | - |
| 70 | F/33 | Ulcerative colitis | Descending colon | + |
| 79 | M67 | Ulcerative colitis | Sigmoid colon | + |
| 80 | M67 | Ulcerative colitis | Descending colon | - |
| 84 | M/30 | Ulcerative colitis | Sigmoid colon | + |
| 85 | M/30 | Ulcerative colitis | Descending colon | - |
| 90 | M/74 | Ulcerative colitis | Sigmoid colon | + |
| 91 | M/74 | Ulcerative colitis | Transverse colon | - |
| 98 | M/57 | Ulcerative colitis | Descending colon | - |
| 99 | M/57 | Ulcerative colitis | Appendiceal orifice | + |
| 104 | M/29 | Ulcerative colitis | Rectum | + |
| 105 | M/29 | Ulcerative colitis | Transverse colon | - |
| 106 | F/33 | Ulcerative colitis | Ascending colon | + |
| 107 | F/33 | Ulcerative colitis | Descending colon | - |
| 108 | M/71 | Ulcerative colitis | Rectum | + |
| 109 | M/71 | Ulcerative colitis | Sigmoid colon | - |
| 110 | M/24 | Ulcerative colitis | Ascending colon | - |
| 111 | M/24 | Ulcerative colitis | Sigmoid colon | + |
| 114 | F/32 | Ulcerative colitis | Sigmoid colon | |
| 115 | F/32 | Ulcerative colitis | Descending colon | + |
| 116 | F/59 | Ulcerative colitis | Sigmoid colon | + |
| 117 | F/59 | Ulcerative colitis | Sigmoid colon | - |
| 118 | F/79 | Ulcerative colitis | Sigmoid colon | + |
| 119 | F/79 | Ulcerative colitis | Rectum | - |
| 120 | M/71 | Ulcerative colitis | Rectum | - |
| 121 | M/71 | Ulcerative colitis | Sigmoid colon | + |
| 122 | M/45 | Ulcerative colitis | Sigmoid colon | + |
| 123 | M/45 | Ulcerative colitis | Transverse colon | - |
| 124 | F/50 | Ulcerative colitis | Rectum | + |
| 125 | F/50 | Ulcerative colitis | Transverse colon | - |
| 126 | F/35 | Ulcerative colitis | Descending colon | + |
| 127 | F/35 | Ulcerative colitis | Sigmoid colon | - |
| 131 | M/28 | Ulcerative colitis | Ascending colon | - |
| 132 | M/28 | Ulcerative colitis | Descending colon | + |

* All samples were paired and collected from the site indicated with and without inflammation in the same patient except sample 19 and 22.

| Sample ID | Gender/Age | HT/PT | Sampling site | Inflammation (-/+) |
|---|---|---|---|---|
| 29 | M/74 | HT | Sigmoid colon | - |
| 40 | F/67 | HT | Sigmoid colon | - |
| 41 | M/74 | HT | Descending colon | - |
| 44 | F/62 | HT | Sigmoid colon | - |
| 45 | M/42 | HT | Sigmoid colon | - |
| 49 | F/56 | HT | Descending colon | - |
| 72 | F/57 | HT | Descending colon | - |
| 73 | F/52 | HT | Descending colon | - |
| 128 | F/53 | HT | Ascending colon | - |
| 129 | M/73 | HT | Ascending colon | - |
| 130 | F/65 | HT | Descending colon | - |
