## Supplemental Figure S1, Table S1, Table S2 for "Live *Malassezia* strains isolated from the mucosa of patients with ulcerative colitis": Table S2. Live fungal strains obtained in the current study.pdf

| Phylum | PT |
|---|---|
| Ascomycota | *Candida albicans* |
| | *Candida glabrata* |
| | *Candida tropicalis* |
| | *Candida orthopsilosis* |
| | *Candida bracarensis* |
| | *Candida parapsilosis* |
| | *Cyberlindnera fabianii* |
| | *Pichia kluyveri* |
| | *Pichia guilliermondii* |
| | *Aspergillus niger* |
| | *Aspergillus unguis* |
| | *Meyerozyma carpophila* |
| | *Meyerozyma guilliermondii* |
| | *Saccharomyces cerevisiae* |
| | *Wickerhamomyces anomalus* |
| | *Kazachstania servazzii* |
| | *Kazachstania aerobia* |
| | *Coniochaeta hoffmannii* |
| | *Clavispora lusitaniae* |
| Basidiomycota | *Malassezia furfur* |
| | *Malassezia obtusa* |
| | *Malassezia globosa* |
| | *Daedaleopsis confragosa* |
| | *Trametes versicolor* |
| | *Rhodotorula mucilaginosa* |
| | *Cryptococcus diffluens* |
| | *Rhodosporidiobolus sp.* |
| Mucoromycota | *Rhizomucor pusillus* |

| Phylum | HT |
|---|---|
| Ascomycota | *Candida albicans* |
| | *Candida glabrata* |
| | *Candida tropicalis* |
| | *Candida orthopsilosis* |
| | *Candida parapsilosis* |
| | *Cyberlinera fabianii* |
| Basidiomycota | *Malassezia sympodialis* |
